# Malaria vaccine candidates displayed on novel virus-like particles are immunogenic and induce transmission-blocking activity

**DOI:** 10.1101/597831

**Authors:** Jo-Anne Chan, David Wetzel, Linda Reiling, Kazutoyo Miura, Damien Drew, Paul R Gilson, David A Anderson, Jack S Richards, Carole A Long, Manfred Suckow, Volker Jenzelewski, Takafumi Tsuboi, Michelle J Boyle, Michael Piontek, James G Beeson

**Affiliations:** Burnet Institute, Melbourne, VIC, Australia; Department of Immunology, Central Clinical School, Monash University, VIC, Australia; ARTES Biotechnology GmbH, Langenfeld, Germany; Technical University of Dortmund, Laboratory of Plant and Process Design, Dortmund, Germany; Laboratory of Malaria and Vector Research, National Institute of Allergy and Infectious Disease, National Institutes of Health, Rockville, Maryland, USA; Department of Medicine, University of Melbourne, VIC, Australia; Proteo-Science Centre, Ehime University, Matsuyama, Ehime, Japan; QIMR-Berghofer Medical Research Institute, Herston, QLD, Australia

## Abstract

The development of effective malaria vaccines remains a global health priority. Currently, the most advanced vaccine, known as RTS,S, has only shown modest efficacy in clinical trials. Thus, the development of more efficacious vaccines by improving the formulation of RTS,S for increased efficacy or to interrupt malaria transmission are urgently needed. The RTS,S vaccine is based on the presentation of a fragment of the sporozoite antigen on the surface of virus-like particles (VLPs) based on human hepatitis B virus (HBV). In this study, we have developed and evaluated a novel VLP platform based on duck HBV (known as Metavax) for malaria vaccine development. This platform can incorporate large and complex proteins into VLPs and is produced in a *Hansenula* cell line compatible with cGMP vaccine production. Here, we have established the expression of leading *P. falciparum* malaria vaccine candidates as VLPs. This includes Pfs230 and Pfs25, which are candidate transmission-blocking vaccine antigens. We demonstrated that the VLPs effectively induce antibodies to malaria vaccine candidates with minimal induction of antibodies to the duck-HBV scaffold antigen. Antibodies to Pfs230 also recognised native protein on the surface of gametocytes, and antibodies to both Pfs230 and Pfs25 demonstrated transmission-reducing activity in standard membrane feeding assays. These results establish the potential utility of this VLP platform for malaria vaccines, which may be suitable for the development of multi-component vaccines that achieve high vaccine efficacy and transmission-blocking immunity.

## INTRODUCTION

Mortality caused by *Plasmodium falciparum* malaria is estimated at 216 million cases annually, with approximately 500,000 deaths occurring worldwide [1]. Despite on-going efforts, malaria control has stalled with little reduction of malaria cases observed in the past few years [1]. The spread of anti-malarial drug resistance together with insecticide resistance in parasite vectors has further escalated the need for an effective malaria vaccine. Malaria vaccine strategies can be broadly classified into three approaches; pre-erythrocytic vaccines that target sporozoites and/or infected hepatocytes, blood-stage vaccines that generally target merozoites and antigens on the surface of infected red blood cells, and transmission-blocking vaccines that target the sexual stages of malaria or mosquito-stage antigens [2]. While vaccines targeting pre-erythrocytic stages and blood stages aim to directly prevent infection and disease, there is a growing focus on vaccines that can interrupt or reduce malaria transmission, highlighted by key global organisations including the World Health Organisation (WHO), Bill and Melinda Gates Foundation and PATH Malaria Vaccine Initiative [3]. The most advanced *P. falciparum* vaccine RTS,S (Mosquirix™) is based on the pre-erythrocytic stage of the parasite life cycle and is the only malaria vaccine to have completed phase III clinical trials [4] and is currently undergoing implementation trials in several African countries [1]. However, vaccine efficacy was low in young children [5] and antibodies induced by vaccination waned quickly in the year after immunisation [6]. The WHO and their partners have set an objective of developing a malaria vaccine with 75% efficacy [3]. To achieve this goal, future vaccine development may be dependent on novel strategies that induce sufficiently high levels of functional antibodies[2].

Transmission-blocking vaccines will need to induce a potent antibody response within the host to inhibit the downstream development of parasites in the mosquito vector after a blood meal [7]. This will prevent or reduce the subsequent spread of malaria parasites throughout an endemic population. However, the advancement of such vaccines remains hampered by the lack of knowledge and tools required to study the sexual, transmissible stages of *P. falciparum*. The progress of sexual-stage antigens as vaccine candidates is further limited, in part, by the difficulty to express high yields of properly folded recombinant antigens, and the requirement for vaccine approaches that generate high and sustained levels of antibodies for effective transmission-blocking activity. Leading vaccine candidates that are expressed during the *P. falciparum* sexual-stage include Pfs230 and Pfs25. Pfs230 is expressed on the surface of gametocytes that reside within the human host, while Pfs25 is expressed on the surface of female gametes, zygotes and ookinetes in the mosquito vector [8]. Numerous studies have shown that individuals naturally exposed to malaria acquire antibodies that target Pfs230 (reviewed in [9]). Therefore, immunity afforded by vaccines based on Pfs230, and other major antigens expressed on the gametocyte surface, have the added benefit of potential for antibody boosting from natural malaria exposure. Pfs25 represents the only sexual-stage antigen that has completed human clinical trials [10–12], and Pfs230 is currently undergoing clinical trials. Antibodies generated through vaccination with Pfs25 have been reported to inhibit the development of parasites within the mosquito midgut [11–13], demonstrating the potential of vaccines to interrupt the transmission of malaria throughout a human population.

The development of potential vaccines requires the induction of strong and sustained immune responses in humans, which has been challenging to achieve. The classic approach of a subunit vaccine includes the delivery of a recombinant antigen formulated with the appropriate adjuvant (reviewed in [14]). Expression of antigens in virus-like particles (VLPs) have been successfully developed for the hepatitis B virus and human papilloma virus (reviewed in [15]). Key characteristic features of viruses such as their repetitive surface geometry and activation of specific immunological receptors are maintained by VLPs. However, they do not have the ability to undergo gene replication as they lack a viral genome [16]. Therefore, they have been established as a safe and effective vaccine delivery platform for use in humans (reviewed in [15]). Furthermore, the particle size of VLPs (ranging from 20–200nm) is optimal for immunogenicity and uptake by dendritic cells [17,18]. VLP-based vaccines elicit strong humoral and cellular responses, supported by their capacity to cross-link B cell receptors for successful B cell activation [19]. The use of VLPs to display and present antigens to the immune system appeared to generate a stronger cellular and humoral response compared to the soluble antigen alone [20,21]. The only VLP-based transmission-blocking vaccine designed for use in human clinical trials involves a chimeric, non-enveloped VLP expressing Pfs25 fused to the Alfalfa mosaic virus coat protein [22]. However, although the vaccine induced Pfs25-specific antibodies in a clinical trial, there was limited inhibition of parasite transmission to mosquitoes [22], suggesting the need for improved vaccine formulations.

Here, we describe a novel platform that incorporates sexual-stage malaria vaccine candidates, Pfs230 and Pfs25, into a VLP scaffold that is based on the duck hepatitis B virus (DHBV) small surface protein (dS). In contrast to the human HBV platform, the DHBV delivery platform allows the expression of large and complex proteins [16], which are typical of malaria vaccine candidates. In this platform, resulting VLPs contain a lipid envelope that favours the optimal presentation of transmembrane and GPI-anchored proteins. *P. falciparum* sexual-stage antigens, Pfs25 and the domains Pfs230c [23] and Pfs230D1M [24] derived from the full-length Pfs230 protein, were genetically fused to the dS and the resulting fusion proteins were co-expressed with wild-type dS in methylotrophic yeast *Hansenula polymorpha* [16]. The successful display of antigens on the chimeric VLPs (referred to as Pfs25-dS/dS, Pfs230c-dS/dS and Pfs230D1M-dS/dS) was confirmed using Western blot analyses and visualized by electron microscopy and super resolution microscopy (presented in the accompanying manuscript by Wetzel D et al 2019, submitted) [25]. In this manuscript, we report the immunogenicity of these sexual-stage chimeric VLPs through animal immunisations and further characterized their functional antibody response through mosquito feeding assays.

## METHODS

### Production and purification of chimeric VLPs

The production and purification of the three different kinds of chimeric VLPs (Pfs230c-dS/dS, Pfs230D1M-dS/dS and Pfs25-dS/dS) was conducted as previously described [16,25]. Genes encoding the three fusion proteins Pfs230c-dS, Pfs230D1M-dS and Pfs25-dS were designed and codon-optimized for expression in *Hansenula polymorpha* (Genbank ref.: MH142260, MH142261, MH142262). Pfs230c-dS consists of amino acids (aa) 443-1132 (630 aa fragment) from full-length Pfs230 [23] and the shorter variant Pfs230D1M-dS consists of aa 542-736 (194 aa fragment) from full-length Pfs230 [24]. The fusion protein Pfs25-dS is comprised of aa 23-193 (170 aa fragment) of the cysteine-rich Pfs25 protein fused to the dS. Each of the fusion proteins encoding genes were co-expressed with a gene encoding the wildtype dS (Genbank ref: MF510122) in a recombinant *Hansenula*-derived yeast cell line. The purification of the chimeric VLPs was based on a downstream process approved for human VLP vaccine production from yeast [26] including adaptations for the purification of VLPs containing the *P. falciparum* antigens [25]. The relative incorporation of the Pfs230-dS/dS or Pfs25-dS/dS fusion proteins into VLPs, compared to total protein, was estimated by densitometry analysis of Coomassie stained gels of VLPs, as described in the accompanying manuscript [25]. The relative incorporation rates were ~30% for Pfs230c-dS/dS VLPs, ~24% for Pfs230D1M-dS/dS VLPs, and ~3% for Pfs25-dS/dS VLPs.

### Expression of monomeric recombinant proteins

The expression of monomeric recombinant Pfs230D1M was performed in HEK293F cells as previously described [27]. Briefly, a truncated form of Pfs230 containing the first 6-cys domain of Pfs230 has previously been expressed as a monomeric recombinant protein in *P. pastoris* termed Pfs230D1H [24]. To assess vaccine-induced antibodies to Pfs230, we expressed a modified form of monomeric recombinant Pfs230D1H in the mammalian HEK293 cell expression system, which we termed Pfs230D1M. Monomeric recombinant Pfs25 was expressed in a wheatgerm cell-free expression system as previously described [28].

### Measuring antibodies to chimeric VLPs by ELISA

Chimeric VLPs or monomeric recombinant proteins were coated onto Maxisorp microtiter plates (Nunc) at 1μg/ml in PBS and incubated overnight at 4°C. Plates were blocked with 1% casein in PBS (Sigma-Aldrich) for 2h at 37°C before primary antibodies were added (polyclonal mouse anti-Pfs230 or anti-Pfs25 antibodies; or rabbit antibodies against Pfs230 or Pfs25). Secondary HRP-conjugated antibodies (polyclonal goat anti-mouse IgG at 1/1000 or anti-rabbit IgG at 1/2500 from Millipore) were used to detect antibody binding. Colour detection was developed using ABTS or TMB liquid substrate (Sigma-Aldrich), which was subsequently stopped using 1% SDS (for ABTS) or 1M sulphuric acid (for TMB). PBS was used as a negative control and plates were washed thrice using PBS with 0.05% Tween in between antibody incubation steps. The level of antibody binding was measured as optical density at 405nm (for ABTS) or 450nm (for TMB).

### Measuring antibody affinity by ELISA

Antibody affinity was assessed using standard ELISA, with the addition of an antibody dissociation step using increasing concentrations of ammonium thiocynate (in PBS). This was incubated for 20 min at room temperature, following incubation with antibody samples (R1917 and 1918 were tested at 1/800, R 1919 and R1920 were tested at 1/100). Colour detection was developed using ABTS (measured at optical density at 405nm).

### Measuring antibodies to the gametocyte surface by flow cytometry

Mature stage V gametocyte-infected erythrocytes were generated as previously described [27,29] and treated with saponin to permeabilize the infected erythrocyte membrane. Gametocytes were subsequently incubated with whole serum from rabbits immunized with Pfs230c-dS/dS VLPs, followed by an AlexaFluor 488-conjugated donkey anti-rabbit IgG (1/500) with ethidium bromide to distinguish between infected and uninfected erythrocytes (1/1000), with washing between each step. Data was acquired by flow cytometry (FACS Verse, BD Biosciences) and analysed using FlowJo software. Antibody levels are expressed as the geometric mean fluorescence intensity (MFI; arbitrary units).

### Immunofluorescence Microscopy

Thin blood smears of stage V 3D7 gametocyte-infected erythrocytes were fixed in 90% acetone and 10% methanol for 5 min at −20°C as previously described [30]. Briefly, slides were rehydrated in PBS and blocked with 6% BSA for 30 minutes before incubation with primary antibodies (rabbit antibodies were tested at 1/100) for 2h. Slides were subsequently incubated with the secondary AlexaFluor 488-conjugated IgG (1/1000; Thermo Scientific) for 1h. Slides were washed thrice in PBS between antibody incubation steps. Slides were mounted in medium containing DAPI to label the parasite nucleus. Images were collected using a Plan-Apochromat (100×/1.40) oil immersion phase-contrast lens (Carl Zeiss) on an AxioVert 200M microscope (Carl Zeiss) equipped with an AxioCam Mrm camera (Carl Zeiss). Images were processed using Photoshop CS6 (Adobe).

### Rabbit immunisations

Pfs230c-dS/dS and Pfs230D1M-dS/dS VLPs were formulated with equal volumes of sterile Alhydrogel aluminium hydroxide vaccine adjuvant (Brenntag, Denmark) and incubated on the shaker for 5 min prior to immunisations. New Zealand White rabbits were immunised 2 weeks apart with 3 doses of chimeric VLPs. The total amount of protein and adjuvant used in each immunisation is presented in Supplementary Tables S1–S3 for Pfs230c-dS/dS, Pfs230D1M-dS/dS and Pfs25-dS/dS VLPs, respectively. Briefly, rabbits (R1864-R1871) were immunised with two vaccine doses of Pfs230c-dS/dS VLPs formulated with and without Alhydrogel. As standard doses, we used 20 μg or 100 μg to total VLP protein for immunizations; the respective amount of Pfs230c protein is reported in Supplementary Tables S1–S3. Rabbits were immunised with Pfs230D1M-dS/dS VLPs formulated with Freund’s adjuvant (R1917, R1918) or Alhydrogel (R1919, R1920) at a single vaccine dose (of 100 μg to total VLP protein). Rabbits were immunised with Pfs25-dS/dS VLPs formulated with Freund’s adjuvant (R1825, R1826) at a single vaccine dose (of 100 μg to total VLP protein). Animal immunisations and the formulation of Pfs25dS/dS VLPs with Freund’s Adjuvant were conducted by the Antibody Facility at the Walter and Eliza Hall Institute. Rabbits were euthanised with a terminal dose of sodium pentobarbitone administered intravenously. Animal immunisations were approved by the Animal Ethics Committee of the Walter and Eliza Hall Institute, Australia (#2017.018).

### Measuring transmission-blocking activity by standard membrane feeding assays

IgG purification from individual rabbit serum samples was performed using Protein G columns (GE Healthcare) according to the manufacturer instructions and adjusted to a final concentration of 20 mg/ml in PBS. The standardized methodology for performing the standard membrane feeding assays (SMFA) was described previously [31]. Briefly, 16-18 days old gametocyte cultures of the *P. falciparum* NF54 line were mixed with purified test IgG at 7.5 mg/ml, and the final mixture, which contained human complement, was immediately fed to ~50 female *Anopheles stephensi* mosquitoes through a membrane-feeding apparatus. Mosquitoes were kept for 8 days and dissected to enumerate the oocysts in the midgut (n=40 per group for the negative control group, and n=20 for test groups). As the negative controls, total IgG purified from pre-immune rabbit sera were utilised. Only midguts from mosquitoes with any eggs in their ovaries at the time of dissection were analyzed. The human serum and red blood cells used for the SMFA were purchased from Interstate Blood Bank (Memphis, TN).

### Statistical analysis

The best estimate of % inhibition in oocyst density (% transmission-reducing activity, %TRA), the 95% confidence intervals (95%CI), and *p*-values from single or multiple feeds were calculated using a zero-inflated negative binomial random effects model (ZINB model) described previously [32]. All statistical tests were performed in R (version 3.5.1), and *p*-values <0.05 were considered significant.

## RESULTS

### Sexual-stage vaccine candidates are displayed on chimeric VLPs

Chimeric VLPs expressing sexual-stage *P. falciparum* antigens, Pfs230 and Pfs25, were produced and purified as described [25]. Antigens were incorporated into the VLP scaffold through genetic fusion of Pfs230 and Pfs25 with the dS protein. For the expression of the chimeric Pfs230 VLPs, we used 2 different constructs. Due to the large size and complexity of Pfs230, we did not attempt expression of full length Pfs230 in VLPs; instead we expressed two truncated forms based on published constructs that have demonstrated induction of transmission-blocking antibodies when used as recombinant protein vaccines in experimental animals. The first was a truncated construct of Pfs230, termed Pfs230c, which includes the first two 6-cysteine domains of Pfs230 [23], was used to generate fusion proteins for VLP formation. Pfs230c has been previously shown to elicit transmission-blocking antibodies [23]. The expression and purification of Pfs230c-dS VLPs was challenging [25]. Therefore, a shorter construct of Pfs230, termed Pfs230D1M (based on the construct described by [24]) which includes only the first 6-cysteine domain of Pfs230, was designed and fused to the dS antigen. Further, Pfs230D1M is a leading transmission-blocking vaccine candidate currently in clinical trials. Here, we evaluated the display of these sexual-stage antigens on the surface of native VLPs. We used specific antibodies generated against Pfs230 and Pfs25 to detect the expression of these antigens on the chimeric VLPs by ELISA (Fig 1). We found that a Pfs230-specific polyclonal antibody recognised the surface of Pfs230c-dS/dS (Fig 1A) and Pfs230D1M-dS/dS VLPs (Fig 1B). Similarly, a Pfs25-specific monoclonal antibody recognised the surface of Pfs25-dS VLPs (Fig 1C). These results confirm the expression of sexual-stage antigens on the surface of chimeric VLPs. There was no recognition of a plain dS VLP without a Pfs230 or Pfs25 fusion protein (Fig 1D).

**Fig 1.**
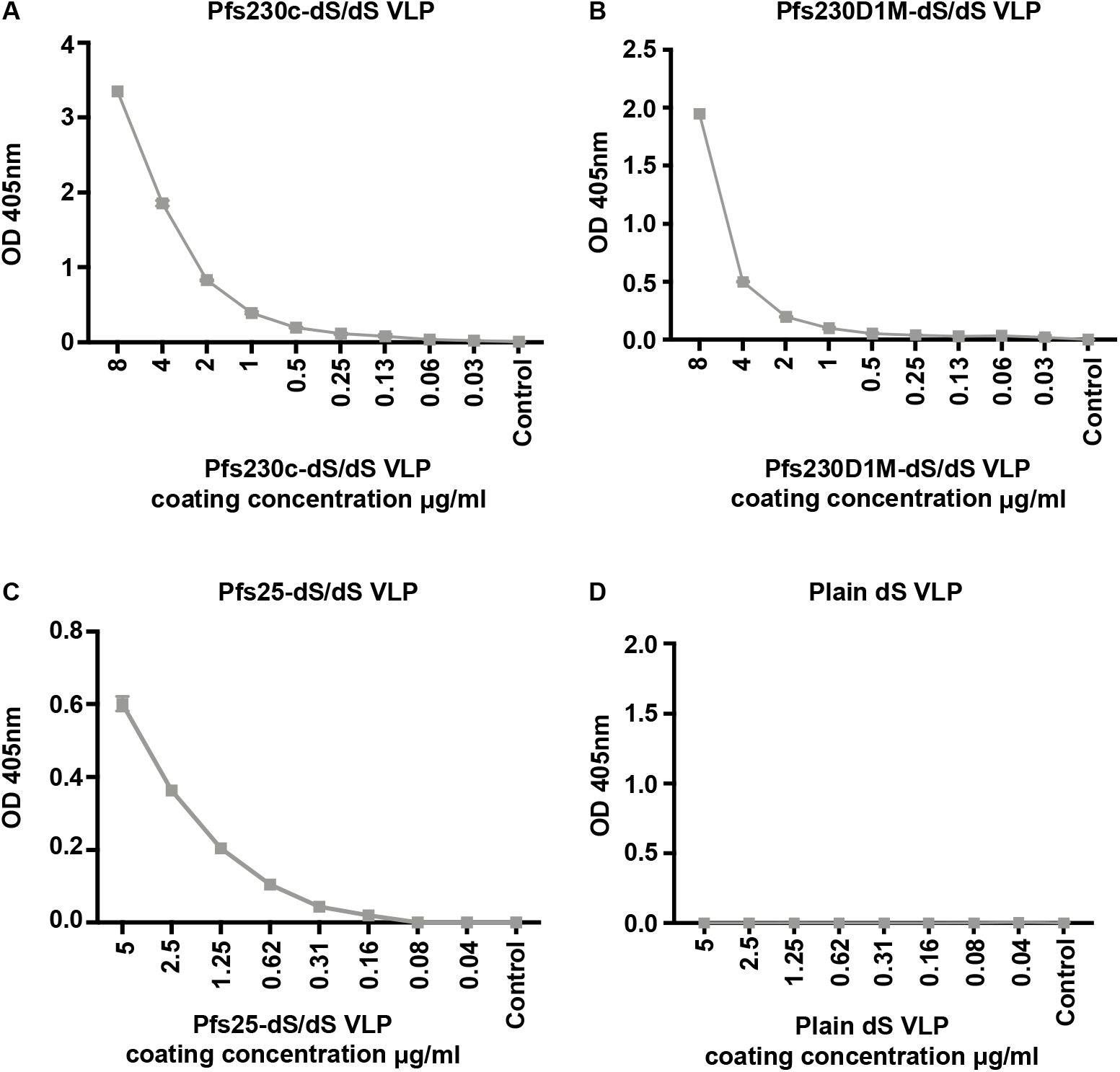
Characterising the expression of sexual-stage antigens on chimeric VLPs. Total antibody binding to (**A**) Pfs230c-dS/dS, (**B**) Pfs230D1M-dS/dS, (**C**) Pfs25-dS/dS and (**D**) plain dS VLPs were measured by ELISA. VLPs were coated at varying concentrations (μg/ml) and probed with either a Pfs25 or Pfs230-specific polyclonal antibody (1μg/ml). In (**D**), plain dS VLPs were probed with anti-Pfs230 antibody. The level of antibody binding is expressed as optical density (OD) measured at 405nm; symbols represent the mean and error bars represent the range between samples tested in duplicate.

### VLPs expressing sexual-stage antigens induced antibodies in rabbits

To understand whether these chimeric VLPs displaying sexual-stage antigens were capable of eliciting an immune response, the VLP constructs were used to immunise rabbits (n=8 for Pfs230c-dS/dS, n=4 for Pfs230D1M-dS/dS, n=2 for Pfs25-dS/dS). Serum from immunised rabbits had significant antibody recognition of monomeric recombinant Pfs230D1M (Fig 2A-D) or Pfs25 (Fig 2E, 2F) in a concentration-dependent manner by ELISA. For immunisations with Pfs230c-dS/dS VLP, we investigated two vaccine doses (20 or 100 μg of total VLP protein) formulated with or without Alhydrogel, which is an adjuvant suitable for clinical use. The higher dose appeared to induce a higher antibody response in rabbits, and formulation with the Alhydrogel adjuvant also contributed to higher antibody levels (Fig 2A, B). Given the findings with Pfs230c-dS/dS VLPs, immunisations with Pfs230D1M-dS/dS VLP were only performed at a single dose (using the higher dose of 100μg of total VLP protein), formulated with Alhydrogel or with Freund’s adjuvant for comparison. Both formulations induced substantial antibodies to monomeric recombinant Pfs230D1M (Fig 2C, D); antibodies were higher for formulations with Freund’s, which is a more potent adjuvant, but cannot be used clinically. For Pfs25-dS/dS VLPs, immunizations were only performed with Freund’s adjuvant as a proof-of-principle, since Pfs25 has already been evaluated with different adjuvants using different platforms in published studies and has advanced into clinical trials [11,13].

**Fig 2.**
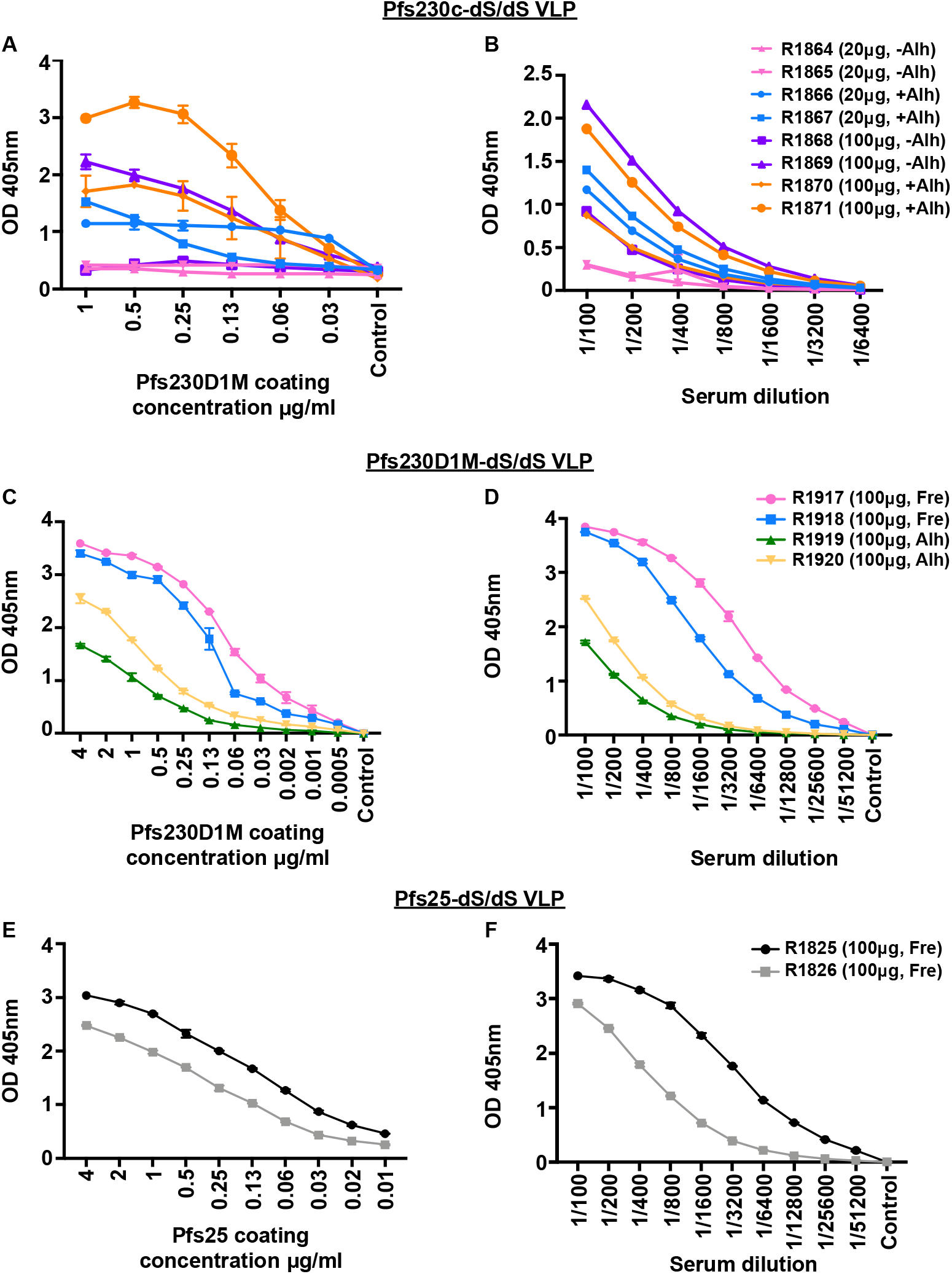
Immunogenicity of sexual-stage chimeric VLPs in rabbits. The level of antibody binding to a titration of monomeric recombinant (**A**, **C**) Pfs230D1M and (**E**) Pfs25 was measured in whole rabbit serum (1/100). Rabbits were immunised with chimeric Pfs230c-dS/dS (**A**, **B**), Pfs230D1M-dS/dS (**C**, **D**) or Pfs25-dS/dS (**E**, **F**) VLPs. Monomeric recombinant proteins were serially diluted from 4μg/ml. The level of antibody binding to monomeric recombinant (**B**, **D**) Pfs230D1M and (**F**) Pfs25 (coated at 1μg/ml) was measured by titrating whole rabbit serum. Rabbits were immunised with chimeric Pfs230c-dS/dS (**A**, **B**), Pfs230D1M-dS/dS (**C**, **D**) or Pfs25-dS/dS (**E**, **F**) VLPs. Whole rabbit serum was serially diluted from 1/100. For all graphs, antibody binding is expressed as optical density (OD) measured at 405nm; symbols represent the mean and error bars represent the range between samples tested in duplicate (n=8 for Pfs230c-dS/dS, n=4 for Pfs230D1M-dS/dS, n=2 for Pfs25-dS/dS). Rabbits R1864-R1871 received Pfs230c-dS/dS VLPs formulated with and without Alhydrogel (**A**, **B**; Alh). Rabbits R1917 and R1918 received Pfs230D1M-dS/dS VLPs formulated with Freund’s adjuvant (Fre), while R1919 and R1920 received Pfs230D1M-dS/dS VLPs formulated with Alhydrogel (**C**, **D**). Rabbits R1825 and R1826 received Pfs25-dS/dS VLPs formulated with Freund’s adjuvant (**E**, **F**). Adjuvants and total VLP protein content used for immunisations are also presented in Supplementary Tables S1–S3.

### Rabbit antibodies induced by chimeric VLPs recognized native Pfs230 expressed on gametocytes

We evaluated whether rabbit antibodies generated against Pfs230c-dS/dS VLPs were capable of recognizing native antigens expressed on the surface of mature stage V gametocytes by flow cytometry (Fig 3A). The majority of the rabbit sera had substantial antibody reactivity to the surface of mature gametocytes, suggesting that vaccine-induced antibodies recognize native Pfs230 (Fig 3A). This finding was confirmed using immunofluorescence microscopy, which showed that rabbit antibodies generated against Pfs230c-dS/dS VLPs labelled native Pfs230 expressed on the surface of fixed mature gametocytes (Fig 3B).

**Fig 3.**
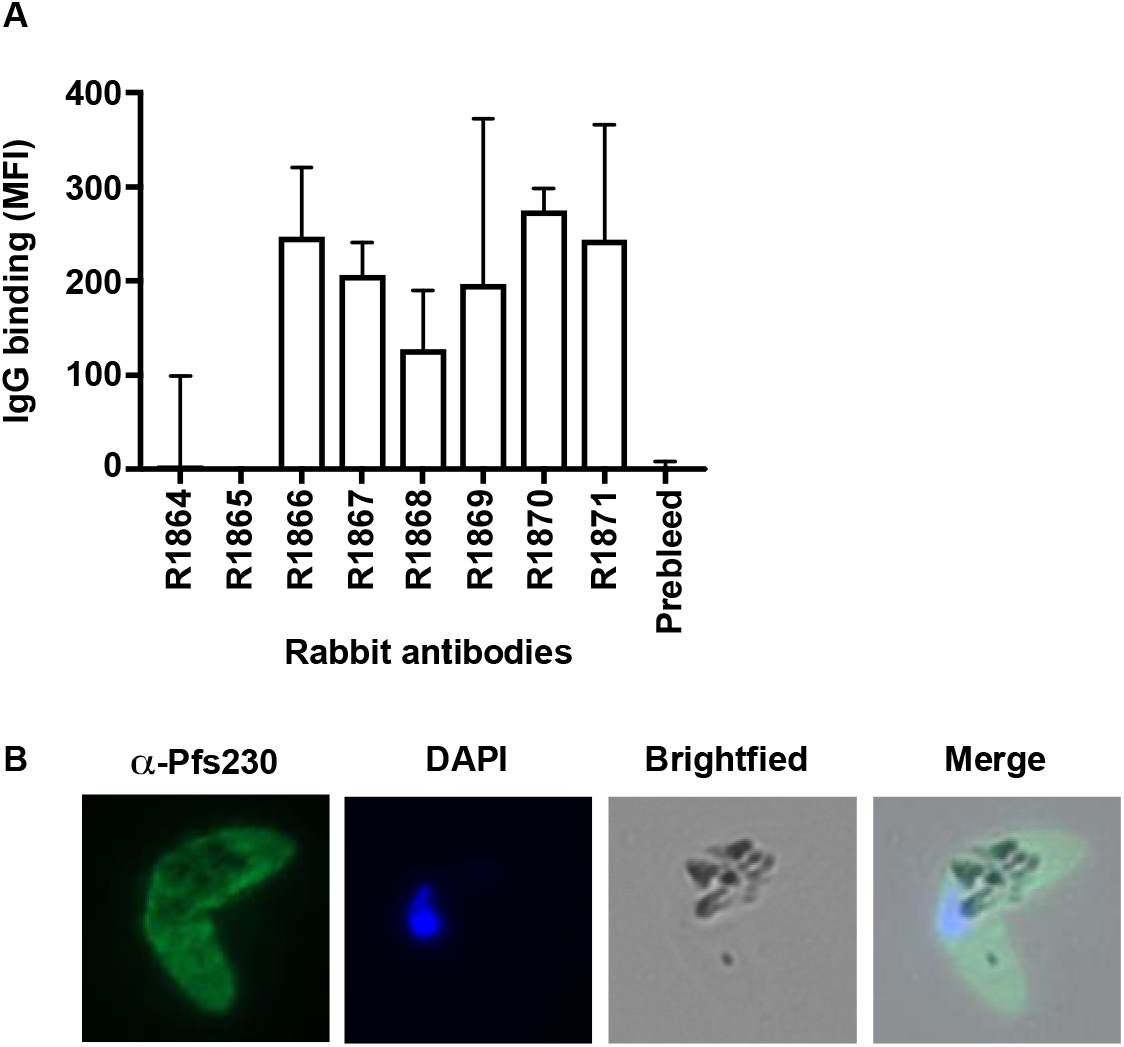
Characterising the antibody function of rabbits immunised with chimeric VLPs. **A**. Total antibody binding to the surface of native *P. falciparum* gametocytes (stage V gametocyte-infected erythrocytes were permeabilised with saponin) measured by flow cytometry. Whole serum from rabbits immunised with Pfs230c-dS/dS VLPs were used (n=8). Antibody levels are expressed as geometric mean fluorescence intensity (MFI); bars represent mean and range of samples tested in duplicate. **B**. Immunofluorescence microscopy demonstrates the recognition of the native gametocyte surface by serum antibodies from rabbits immunised with Pfs230c-dS/dS VLPs (R1870; green). Cells were fixed with 90% acetone and 10% methanol, and DAPI was used to stain nuclear DNA (blue). Representative images taken of gametocytes labelled with R1870 are shown.

### Rabbit antibodies induced by chimeric VLPs have transmission-reducing activity

To address the functional significance of rabbit antibodies generated against sexual-stage chimeric VLPs, we examined the ability of these antibodies to inhibit mosquito infection through standard membrane feeding assays (SMFA) in the presence of human complement [31]. Functional transmission-blocking activity is defined as the reduction in oocyst count compared to a negative control group. When antibodies against Pfs230c-dS/dS VLPs were tested at 7.5mg/ml, none of them showed significant inhibition (Table 1). In contrast, 2 out of 4 rabbits immunized with Pfs230D1M-dS/dS VLPs demonstrated significant inhibition in SMFA (Table 2). Antibodies generated against Pfs25-dS/dS VLPs from one of the two rabbits also successfully blocked the development of oocysts within the mosquito midgut (Table 3; 97.4% inhibition). It was interesting that the second rabbit did not significantly inhibit transmission despite substantial induction of antibodies detected by ELISA, suggesting that antibody titre and specificity may be important for functional activity. Interestingly, there was no clear relationship between IgG levels and activity in SMFA by rabbit antibodies generated to Pfs230D1M-dS/dS. While VLPs formulated with Freund’s adjuvant generated higher IgG reactivity to Pfs230, IgG generated using VLPs formulated with alum tended to have stronger inhibitory activity in SMFA (Table 2). To understand these differences further, we estimated the avidity of IgG binding to Pfs230D1M for different rabbit antibodies (Supplementary Figure S1). This indicated no major difference in IgG avidity that would explain difference in activity in SMFA, suggesting differences in functional activity may be explained by differences in epitope targeting of antibodies.

**Table 1.** Assessment of transmission-blocking activity of rabbit antibodies against Pfs230c-dS/dS VLPs by SMFA

| Rabbit ID<br>(+/- Adjuvant) | Protein content<sup>1</sup> | Antibody<br>levels<sup>2</sup> | %<br>inhibition<sup>3</sup> | 95% confidence<br>intervals | *p* value |
| --- | --- | --- | --- | --- | --- |
| R1864<br>(-Alhydrogel) | 20µg VLP<br>~6µg Pfs230c-dS | 0.29 | -20.9 | -162.2 – 43.5 | 0.62 |
| R1865<br>(-Alhydrogel) | 20µg VLP<br>~6µg Pfs230c-dS | 0.30 | 6 | -100.0 – 57.2 | 0.85 |
| R1866<br>(+Alhydrogel) | 20µg VLP<br>~6µg Pfs230c-dS | 1.17 | -30.8 | -187.4 – 43.5 | 0.52 |
| R1867<br>(+Alhydrogel) | 20µg VLP<br>~6µg Pfs230c-dS | 1.40 | -34.8 | -193.4 – 37.7 | 0.46 |
| R1868<br>(-Alhydrogel) | 100µg VLP<br>~30µg Pfs230c-dS | 0.92 | -13.7 | -143.7 – 48.3 | 0.72 |
| R1869<br>(-Alhydrogel) | 100µg VLP<br>~30µg Pfs230c-dS | 2.16 | -28.9 | -182.5 – 41.8 | 0.53 |
| R1870<br>(+Alhydrogel) | 100µg VLP<br>~30µg Pfs230c-dS | 0.87 | -8.1 | -124.4 – 50.5 | 0.90 |
| R1871<br>(+Alhydrogel) | 100µg VLP<br>~30µg Pfs230c-dS | 1.88 | -16.6 | -156.6 – 48.0 | 0.70 |

<sup>1</sup>Refers to total VLP protein content and estimated incorporation rate of Pfs230c-dS
<sup>2</sup>Mean antibody levels by ELISA at 1/100 serum dilution (OD 405nm)
<sup>3</sup>% inhibition is the best estimate from 1 feed
All rabbit antibodies were tested at an IgG concentration of 7.5 mg/ml.

**Table 2.** Assessment of transmission-blocking activity of rabbit antibodies against Pfs230D1M-dS/dS VLPs by SMFA

| **Rabbit ID**<br>**(Adjuvant)** | **Antibody levels<sup>1</sup>** | **% inhibition<sup>2</sup>** | **95% confidence**<br>**intervals** | **p value** |
| --- | --- | --- | --- | --- |
| R1917<br>(Freund's) | 3.84 | 43.7 | 4.1 – 68.0 | **0.03** |
| R1918<br>(Freund's) | 3.74 | 9.2 | -51.3 – 47.5 | 0.74 |
| R1919<br>(Alhydrogel) | 1.72 | 40.7 | -0.2 – 65.9 | 0.05 |
| R1920<br>(Alhydrogel) | 2.51 | 91.7 | 85.7 – 95.4 | **0.001** |

<sup>1</sup>Mean antibody levels by ELISA at 1/100 serum dilution (OD 405nm)
<sup>2</sup>% inhibition is the best estimate from 2 feeds
Protein content was 100µg to total VLP protein and 24µg Pfs230D1M-dS; all rabbit antibodies were tested at an IgG concentration of 7.5 mg/ml; *p* values that achieved statistical significance are presented in bold.

**Table 3.** Assessment of transmission-blocking activity of rabbit antibodies against Pfs25-dS/dS VLPs by SMFA

| **Rabbit ID**<br>**(Adjuvant)** | **Antibody levels<sup>1</sup>** | **% inhibition<sup>2</sup>** | **95% confidence**<br>**intervals** | **p value** |
| --- | --- | --- | --- | --- |
| R1825<br>(Freund's) | 3.42 | 97.4 | 95.4 – 98.8 | **0.001** |
| R1826<br>(Freund's) | 2.91 | -1.4 | -69.4 – 40.5 | 0.96 |

<sup>1</sup>Mean antibody levels by ELISA at 1/100 serum dilution (OD 405nm)
<sup>2</sup>% inhibition is the best estimate from 2 feeds
Protein content was 100µg to total VLP protein and 3µg Pfs25-dS; all rabbit antibodies were tested at an IgG concentration of 7.5 mg/ml; *p* value that achieved statistical significance is presented in bold.

## DISCUSSION

Our work describes the immunogenicity of transmission-blocking malaria vaccine candidates presented on the surface of VLPs using a novel platform based on the duck hepatitis B virus. Leading sexual-stage antigens, Pfs230 and Pfs25 were engineered into chimeric VLPs and their surface expression characterised using specific antibodies. Data is presented in the accompanying manuscript [25] on the production and purification of these chimeric VLPs. These showed that chimeric VLPs expressed proteins of expected size corresponding to Pfs25 and the respective Pfs230 domains, including the small surface protein dS. Imaging by electron microscopy identified homogenous, particulate structures corresponding to the chimeric VLPs and super resolution microscopy further confirmed the recognition of Pfs230-derived domains and Pfs25 on the chimeric VLP surface, and their colocalisation with the dS antigen, using specific polyclonal antibodies. In this study, we showed that the VLPs were reactive by ELISA with antibodies to Pfs230 or Pfs25, further demonstrating the display of malaria vaccine candidates on the VLP surface. In rabbit immunisation studies, we showed that the VLPs were immunogenic and were capable of inducing substantial antibody reactivity to monomeric recombinant Pfs230D1M or Pfs25 protein. Further, rabbit antibodies generated against Pfs230c-dS/dS and Pfs230D1M-dS/dS VLPs recognised the surface of native gametocytes by flow cytometry and this was also visualised by immunofluorescence microscopy. Importantly, rabbit antibodies against Pfs230D1M-dS/dS and Pfs25-dS/dS VLPs were capable of blocking mosquito infection measured through standard membrane feeding assays.

A conformational-dependent monoclonal antibody, 4B7, was used to characterise the display of antigen expressed by the Pfs25-dS/dS VLP. There were high levels of antibody recognition measured by ELISA to the Pfs25-dS/dS VLP, which correlated with the coating concentration of monomeric recombinant Pfs25. Similarly, using a Pfs230-specific polyclonal antibody, the display of antigen expressed by the Pfs230c-dS/dS and Pfs230D1M-dS/dS VLP was confirmed. This suggests that the Pfs25 and Pfs230 constructs had successfully been incorporated as fusion proteins into the chimeric VLPs. The incorporation rate of Pfs25-dS into VLPs was relatively low and optimization may be needed to increase incorporation rates prior to further evaluation of Pfs25 VLPs using this platform.

Our immunogenicity studies showed that the chimeric VLPs could elicit an immune response in rabbits. For immunisations with Pfs230c-dS/dS VLP, two dosing regimens were used in the presence and absence of Alhydrogel. As the higher dose appeared to induce a better antibody response in rabbits, and formulation with Alhydrogel was needed for high antibody levels, immunisations with Pfs230D1M-dS/dS VLP were done at a single dose (using the higher dose) and formulated with Alhydrogel. Pfs230D1M-dS/dS VLPs were also formulated with Freund’s adjuvant for comparison. VLPs formulated with Alhydrogel induced substantial antibody responses, but responses were higher with Freund’s adjuvant. This was expected because Freund’s is a more potent adjuvant than Alhydrogel. Our results suggest that investigating other clinically-acceptable adjuvants that may induce higher antibody responses than Alhydrogel would be valuable. Further, rabbits were immunised with Pfs25-dS/dS VLP using only Freund’s adjuvant as a proof-of-principle to evaluate immunogenicity and transmission-blocking activity. Antibodies generated against the Pfs25-dS/dS VLP recognised monomeric recombinant Pfs25 by ELISA. Together, these findings suggest that immunisation with chimeric VLPs induced a good immune response in rabbits, further supporting the potential for the use of this VLP platform to present malaria antigens for vaccination. However, further detailed studies are needed to evaluate the immunogenicity and functional activity of antibodies generated by antigens presented as VLPs compared to monomeric recombinant proteins. In addition, we measured the ability of rabbit antibodies to recognise the surface of native gametocytes by flow cytometry. We found that rabbit antibodies generated against Pfs230c-dS/dS VLPs were capable of recognising the surface of mature, stage V gametocytes. This was also confirmed by immunofluorescence microscopy using fixed smears of mature gametocytes.

Functional antibody responses were evaluated using standard membrane feeding assays, which are widely used to assess transmission-reducing activity [32]. Antibodies from rabbits immunised with Pfs25-dS/dS VLPs and Pfs230D1M-dS/dS could strongly inhibit the development of oocysts within the mosquito midgut, indicating the successful inhibition of parasite transmission. However, inhibitory activity was variable between immunized rabbits. While further studies with larger numbers of animals, and including mice or rats, will be required to better understand variability in vaccine responses, the findings here are sufficient to establish a proof-of-concept for generating transmission-blocking activity using VLPs. Our findings support previous work that showed that rabbit antibodies against Pfs230D1M had transmission-reducing activity by SMFA [24]. Recent work has also reported that only constructs containing domain 1 of Pfs230 were capable of inducing transmission-reducing activity by SMFA, compared to constructs lacking that particular domain [33].

Further, the Pfs230D1M construct is currently undergoing phase I clinical trials. Together, these findings support the importance of Pfs230D1M in the induction of functional transmission-reducing antibodies. Of note, transmission-reducing activity was not observed for antibodies generated against Pfs230c-dS/dS VLPs. This could have been due to the sub-optimal folding of Pfs230c, such that key functional epitopes were not displayed, or important antibody-binding epitopes were masked. These factors potentially resulted in low purity and reduced yield of chimeric VLP and thus led to the lack of functional antibodies generated. Interestingly, there was no clear relationship between IgG reactivity levels quantified by ELISA and transmission-blocking activity in SMFA. This was particularly evident for Pfs230D1M-dS/dS; immunization with VLPs formulated with Freund’s adjuvant generated higher IgG reactivity than using alum as the adjuvant, but IgG generated using alum tended to have greater activity in SMFA. There was no substantial difference in avidity between IgG induced using different adjuvants, which suggests the differences in transmission-blocking activity are most likely explained by differences in epitope-specificity of IgG. Future studies with a larger number of animals and immunization regimens may help further understand this.

In conclusion, we have demonstrated the successful display of sexual-stage antigens Pfs230 and Pfs25 on the surface of chimeric VLPs. We have established a proof-of-concept which showed that these VLPs generated significant immune responses that recognised homologous recombinant protein and native sexual-stage antigens expressed on gametocytes. Further, these antibodies had the ability to block the transmission of parasites to mosquitoes through membrane feeding assays. Together, our results support the further evaluation of chimeric VLPs as a novel delivery platform for leading malaria vaccine candidates. Future studies to optimise antigen incorporation and presentation in VLPs and to evaluate different adjuvants and dosing regimens for immunization will further inform the potential utility of this strategy for malaria vaccine development.

## Author contributions statement

Conceptualization, JC, DW, LR, PG, JR, DA, MS, MP, and JB; Methodology, JC, DW, LR, DD, VJ, KM, TT and MP; Investigation, JC, DW, LR, DD and KM; Administration, VJ, JR, DA, MP and JB; Manuscript preparation, JC, DW and JB (and critically reviewed by all authors); Funding Acquisition, JB and MP. Supervision, JB, MP, CL, MB. All authors approved the final manuscript.

## Acknowledgements

The authors gratefully acknowledge Colleen Woods for ongoing advice and project feedback, Heribert Helgers, Renske Klassen, Christine Langer, Ashley Lisboa-Pinto and Thomas Rohr for technical and academic assistance.

## Funding

Funding was provided by PATH Malaria Vaccine Initiative and the National Health and Medical Research Council (NHMRC) of Australia (Senior Research Fellowship and Program Grant to JB, Career Development Fellowship to MB). Burnet Institute is supported by funding from the NHMRC Independent Research Institutes Infrastructure Support Scheme and a Victorian State Government Operational Infrastructure grant. The SMFA work performed here was supported was supported in part by the intramural program of the National Institute of Allergy and Infectious Diseases/NIH. The funders had no role in study design, data collection and analysis, decision to publish or preparation of the manuscript. ARTES Biotechnology GmbH provided support in the form of salaries for authors (DW, MS, VJ and MP) and generation of VLPs, but did not have any additional role in the study design, data collection and analysis, decision to publish or preparation of the manuscript. The specific roles of these authors are articulated in the ‘Author Contributions’ section.

## Competing Interests

The authors DW, MS, VJ and MP are associated with ARTES Biotechnology GmbH which owns the license for the VLP technology [34,35]. This does not alter our adherence to PLOS ONE policies on sharing data and materials.

## SUPPLEMENTARY MATERIALS

**Supplementary Table S1.**
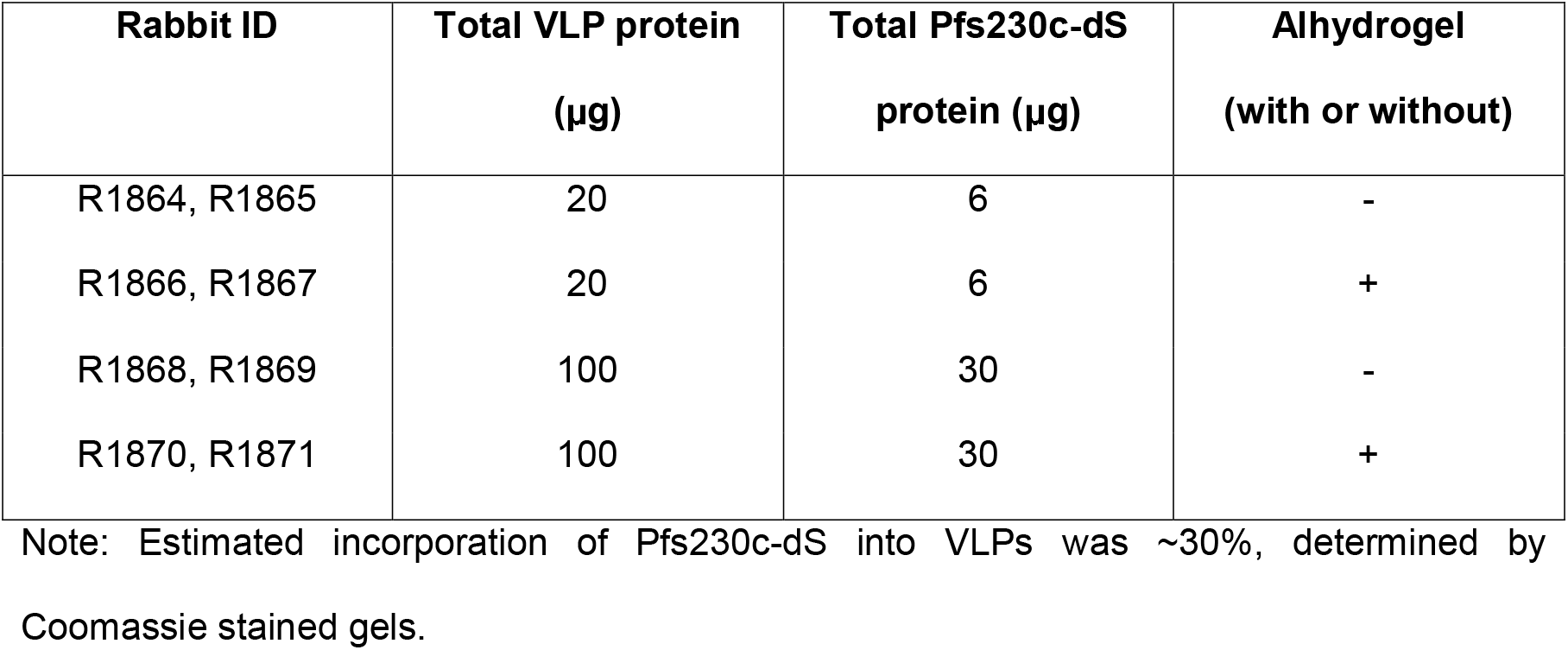
Vaccine groups for Pfs230c-dS/dS VLP rabbit immunisations

| **Rabbit ID** | **Total VLP protein**<br>**(µg)** | **Total Pfs230c-dS**<br>**protein (µg)** | **Alhydrogel**<br>**(with or without)** |
| --- | --- | --- | --- |
| R1864, R1865 | 20 | 6 | - |
| R1866, R1867 | 20 | 6 | + |
| R1868, R1869 | 100 | 30 | - |
| R1870, R1871 | 100 | 30 | + |

Note: Estimated incorporation of Pfs230c-dS into VLPs was ~30%, determined by Coomassie stained gels.

**Supplementary Table S2.** Vaccine groups for Pfs230D1M-dS/dS VLP rabbit immunisations

| **Rabbit ID** | **Total VLP protein**<br>**(µg)** | **Total Pfs230D1M-**<br>**dS protein (µg)** | **Adjuvant** |
| --- | --- | --- | --- |
| R1917, R1918 | 100 | 24 | Freund's |
| R1919, R1920 | 100 | 24 | Alhydrogel |

Note: Estimated incorporation of Pfs230D1M-dS into VLPs was ~24%, determined by Coomassie stained gels.

**Supplementary Table S3.** Vaccine groups for Pfs25-dS/dS VLP rabbit immunisations

| **Rabbit ID** | **Total VLP protein**<br>**(µg)** | **Total Pfs25-dS**<br>**protein (µg)** | **Adjuvant** |
| --- | --- | --- | --- |
| R1825, R1826 | 100 | 3 | Freund's |

Note: Estimated incorporation of Pfs25-dS into VLPs was ~3%, determined by Coomassie stained gels.

**Fig S1.**
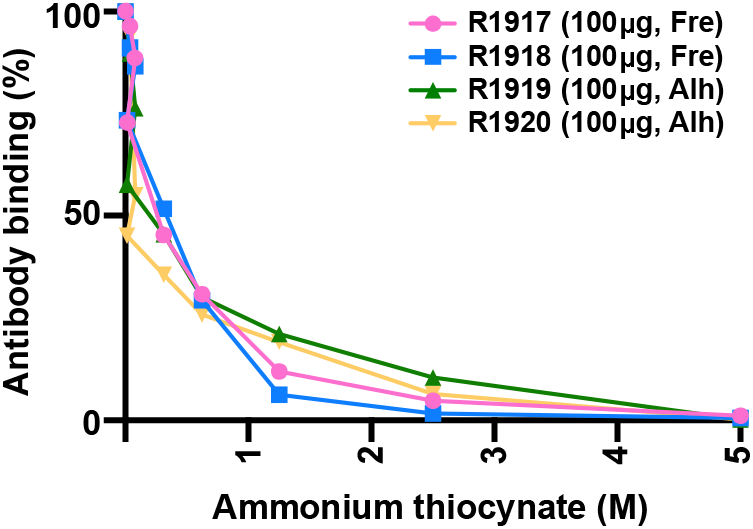
Avidity of rabbit antibodies to bind monomeric recombinant Pfs230D1M. The avidity of antibody binding to monomeric recombinant Pfs230D1M was assessed in serum from rabbits immunised with Pfs230D1M-dS/dS VLPs. The affinity of rabbit antibodies to bind monomeric recombinant Pfs230D1M was similar, irrespective of adjuvant formulation used for rabbit immunisations. Rabbits R1917 and R1918 received Pfs230D1M-dS/dS VLPs formulated with Freund’s adjuvant (Fre), while R1919 and R1920 received Pfs230D1M-dS/dS VLPs formulated with Alhydrogel (Alh). Antibody affinity was assessed by the dissociation of antibodies using increasing concentrations of ammonium thiocynate. Antibody binding is defined as the OD of ammonium thiocynate-treated samples/OD of untreated samples x100.

